# Uncertainty in cell fate decision making: Lessons from potential landscapes of bifurcation systems

**DOI:** 10.1101/2021.01.03.425143

**Authors:** Anissa Guillemin, Elisabeth Roesch, Michael P.H. Stumpf

## Abstract

Cell fate decision making is known to be a complex process and is still far from being understood. The intrinsic complexity, but also features such as molecular noise represent challenges for modelling these systems. Waddington’s epigenetic landscape has become the overriding metaphor for developmental processes: it both serves as pictorial representation, and can be related to mathematical models. In this work we investigate how the landscape is affected by noise in the underlying system. Specifically, we focus on those systems where minor changes in the parameters cause major changes in the stability properties of the system, especially bifurcations. We analyse and quantify the changes in the landscape’s shape as the effects of noise increase. We find ample evidence for intricate interplay between noise and dynamics which can lead to qualitative change in a system’s dynamics and hence the corresponding landscape. In particular, we find that the effects can be most pronounced in the vicinity of the bifurcation point of the underlying deterministic dynamical systems, which would correspond to the cell fate decision event in cellular differentiation processes.

## 1 Introduction

Cell-fate decision making processes are central to developmental biology. These apparently highly choreographed processes are responsible for cells to change into new, well defined states. Many facets of cell fate decision making have been investigated, and improved experimental technologies allow us to investigate these fundamental events in great detail. Theoretical studies are slowly catching and increasingly allow us to make sense of the available and emerging data.

Waddington’s epigenetic landscape is the most famous metaphor used in this context [1]: and it has inspired both the experimental investigation, as well as the classical and modern theoretical investigations into cell-fate decision making: a cell is rolling from the top of a mountain into the bottom of a valley, following the contours of the valleys and hills and choosing its paths among all possible. In this simplified illustration, the three dimensions of Waddington’s epigenetic landscape have very specific interpretations. First, the *x*-axis represents the phenotype, *e.g.* defined by the levels of expression of the underlying genes. A stable gene expression pattern is called a cell type and all of them are separated along this axis. Second, the y-axis represents the time-dependent dimension in which the process occurs. It starts with the initial condition and ends when a cells’ final gene expression pattern attains a stable state (a valley bottom). One can identify this dimension with the system’s driver [2]. And finally, the *z*-axis corresponds to the system’s potential [3,4], which via its shape determined the dynamics associated with the developmental process. Often, and indeed originally, this has been likened to potential energy surfaces or the energy needed for browsing the landscape from one point to another, as the image of the gravitational forces. This potential is governed by a set of gene regulatory elements – DNA, mRNAs, proteins and small molecules – that in concert control the cell fate [5]. From this point of view, qualitative changes of the landscape’s shape will modify the cellular dynamics of fate decisions. In order to conserve the cellular population balance in a multicellular system, the gene regulatory network (GRN) governing the shape of the potential landscape must operate within defined limits; otherwise changes in the landscape will distort the differentiation dynamics, potentially with detrimental outcomes. However, within the GRN molecular noise does exist. Here we denote this noise as the transcriptional stochasticity [6,7]. For cells this can allow considerable levels of flexibility in the fate decision making process. Unfortunately, however, for non-linear dynamical systems the interplay between non-linearities in the dynamics (for example, as encoded in the landscape) and noise largely defies intuition [8] and makes the ensuing processes challenging to describe or understand. For even very simple systems we can observe complex dynamics, once stochasticity and non-linear dynamics both influence a system’s dynamics.

Inspired by the emerging theory of dynamical systems, Waddington’s representation has from the outset enabled and encouraged the application to link mathematical descriptions to cell fate decision making processes [9]. In particular it has encouraged the analysis of development in terms of bifurcations: qualitative changes in a system’s dynamics as parameters are varied. Bifurcations and their analysis have been central to the analysis of deterministic dynamical systems, in particular, ordinary differential equations (ODE). Despite important foundational work in this area dating back to at least the 1970s, much less is known about qualitative changes, their determinants, and their ramifications in stochastic dynamical systems. The extent to which the bifurcation structure of an ODE is reflected in the dynamics of, *e.g.* a stochastic differential equation (SDE) varies, depending on the system dynamics and the type of noise [10,11]. Here we explore this interplay for a set of generic bifurcations (for deterministic dynamics) and a well-defined and tunable choice for noise. Noise is incorporated via SDEs (see below) and we use simulations to construct the (empirical) Waddington landscapes corresponding to the different types of bifurcations, and for varying strengths of noise. Thus, a valley in the landscape corresponds to a stable fixed point in the differential equation system and a hill to an unstable fixed point (well defined in Section 2.1). The point where the system qualitatively changes is called the bifurcation event and, in our case represents the commitment to a specific cell fate. We systematically explore how this landscape changes upon the introduction of noise. These associations between specific bifurcations and biological processes have already been used in the past [2, 12–15]. For instance, a cell differentiating into two different lineages (potential fates) is often seen to be represented by a supercritical pitchfork bifurcation: the split of a valley into two distinct valleys in the epigenetic landscape point of view [2].

Here we focus on the canonical co-dimension 1 bifurcations [16] and construct their empirical quasi-potential landscapes. These bifurcations are chosen because they represent many important cell-fate decision making systems in simplified form. We then use these to explore how qualitative and quantitative features, including measures of variability change with noise. We note, that both molecular noise and (deterministic) qualitative change in cell state have been explored in some detail in the biological

literature. There has been, however, a lack of analyses, where both noise and nontrivial dynamics are simultaneously considered. As hinted at above, there are major conceptual challenges even at the purely mathematical level in any such analysis.

## 2 Methods

### 2.1 Stochastic differential equations

Differential equations provide a convenient and popular framework to model many dynamical systems in biology [11,17]. Ordinary differential equations (ODE) are used to describe continuous deterministic dynamical systems, and stochastic differential equations (SDE) are one way to study the corresponding stochastic dynamics. Formally, we can write for the ODE

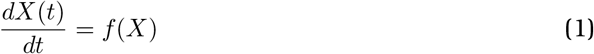

where *X* is the (potentially vector-valued) state of the system, *t* denotes time, and *f* (*X*) describes the deterministic dynamics of the system.

For SDEs we have to replace the derivative by differentials and write

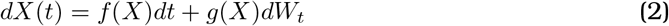

where again *X* represents the system state, *t* represents the time, *f* (*X*) – also referred to as the drift term – describes the deterministic part of the dynamics [18], and *g*(*X*) – also referred to as the diffusion term – captures the stochastic aspects of the dynamics. *dW_t_* is a Wiener process increment (see [18,19] for detailed discussions). In *g*(*X*) the type of the scaling of the noise is defined; we implement geometric random noise, where the diffusion term of the SDE depends linearly on the system state

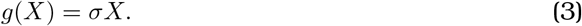

Other choices are possible, but the advantage of this type of noise is that it gives rise to sufficiently non-trivial behaviour: in particular the distributions of X for the simple *geometric Brownian motion process,* given by *dX* = *μXdt* + *σXdW_t_*, can be shown to be non log-Normal, *i.e.* non-Gaussian.

### 2.2 Co-dimension 1 bifurcations

A bifurcation in a deterministic dynamical system is associated with qualitative change in the stability of the stationary solutions [16]. In this work we focus on co-dimension 1 bifurcations [11]; they differentiate themselves from other bifurcations as in that the qualitative stability changes in the systems are caused by a single bifurcation parameter (instead of a set of parameters). We here consider the three canonical co-dimension 1 bifurcations in their simplest normal forms given by

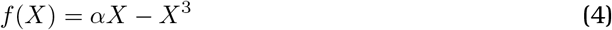

for the *supercritical pitchfork* bifurcation,

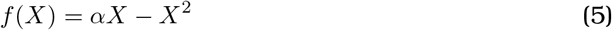

for the *transcritical* bifurcation and

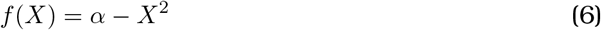

for the *saddle-node* bifurcation. We consider SDEs with drift terms given by these equations, and diffusion terms corresponding to geometric random noise.

### 2.3 Potential landscapes and uncertainty measure

We estimate the Waddington’s epigenetic landscape for a system with the states X using the quasi-potential function,

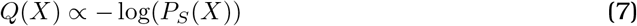

where *P_S_* is a the probability density function of the observed steady state distribution. *P_S_* is obtained through simulations, as described elsewhere [4,13,20].

A simplified example for the construction of *Q*(*X*) is given in Figure 1 where a range of dynamical systems is defined for exemplar values of the bifurcation parameter *α* and two noise levels. We constrain *α* to the interval [-4,4] and the time series data together with the corresponding quasi-potential function are displayed for scenarios with and without noise. To construct the potentials (via the stationary distribution) we solve the dynamics for a set of different initial conditions, *IC*. From this, *P_s_* (*X*) is estimated using kernel density estimation (as described elsewhere).

**Figure 1:**
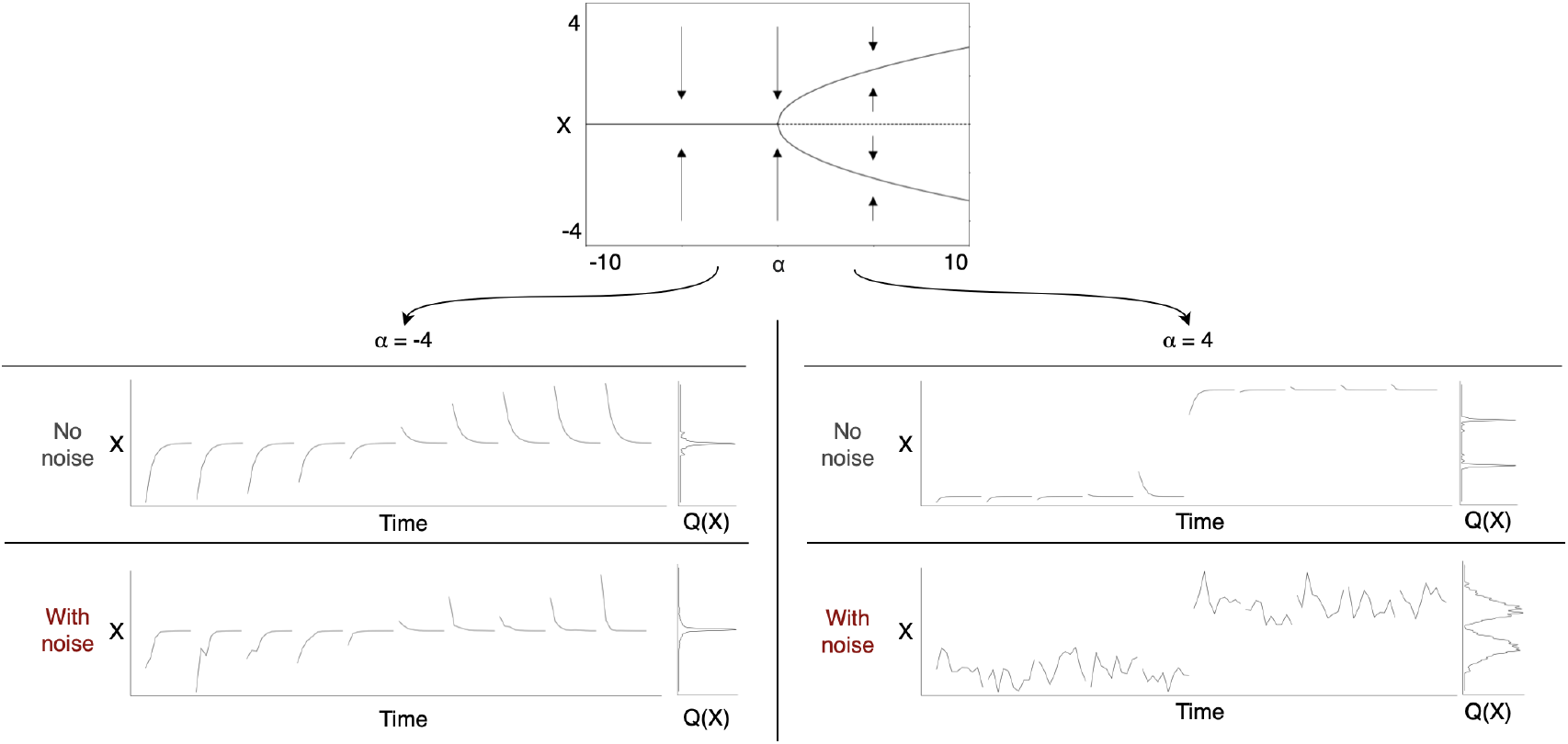
Top: The bifurcation diagram of the supercritical pitchfork bifurcation where stable (solid line) and unstable (dashed line) fixed points of the system are shown as a function of the bifurcation parameter α. Left: For negative α, the two time courses with no noise (deterministic setting) and simulations with noise (stochastic setting) are shown. They both contain trajectories of ten initial conditions. Appended to the time course plot is the quasi-potential function built based on the ten simulations. In both cases (no noise and with noise) we observe a clear peak in the graph of the Junction at X = 0. The peak aligns with the stable fixed point at X = 0 for negative a in the supercritical pitchfork bifurcation. Right: Again, the deterministic and stochastic cases are shown but for positive values of α. The graphs of the quasi-potential function peaks at the positions of the stable fixed points at 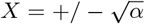.

We use the entropy,

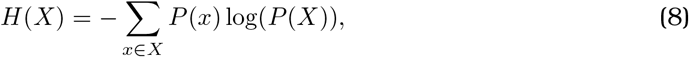

to quantify variability in system states as dynamics chance upon the introduction of noise.

## 3 Results

### 3.1 Quasi-potential landscape of the supercritical pitchfork bifurcation for increasing noise

We first investigate the quasi-potential for the supercritical pitchfork bifurcation, to observe if and how the epigenetic landscape is qualitatively affected by noise. As shown in Figure 2.a, we observe valleys in the empirical quasi-potential function where stable fixed points are expected for the deterministic system (at *X* = 0 for negative α and 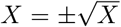 for positive α). The quasi-potential thus reflects qualitatively aspects of the bifurcations of the deterministic dynamics. Introduction of geometric noise, Equation (2), alters the shape of the empirical quasi-potentials shown in Figure 2.b–d, and sometimes quite profoundly: while for *α* < 0 we identify a clear valley at all noise levels, for *α* > 0 valleys become shallower and for larger levels of noise they become flat to the point of merging and/or disappearing altogether.

**Figure 2:**
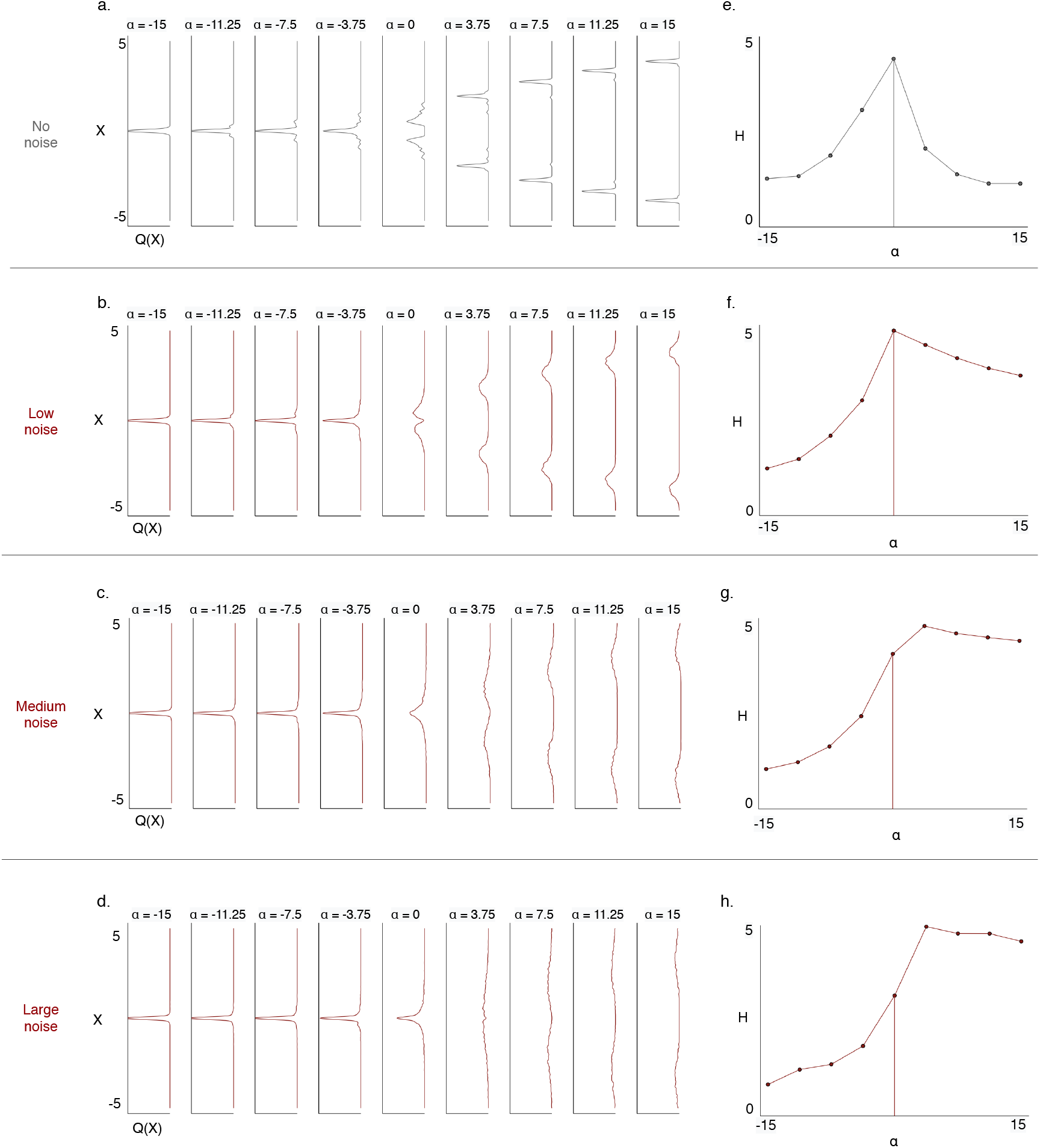
The quasi-potential Q(X) (a-d) and the entropy over the state space H(X) (e-h) are visualised as a system undergoes a supercritical pitchfork bifurcation for different representative noise levels (0.0 < *σ* < 1.6). The first row (a and e) represents a system with no noise (*σ* = 0.0); here the steady state probability distribution in state space would (for *t* → ∞) be a set of Dirac *δ*-functions (the invariant set of the dynamics [8,21]); for convenience we have have stuck to finite time-series and hence we observe broadened peaks. The next three rows show the same landscapes and entropies but for noisy dynamics with *σ* = 0.6 in b and f, *σ* = 1.3 in c and g, and *σ* = 1.6 in d and h (which reflects also observations for larger noise levels).

We can further quantify these changes by calculating the entropy of the state variable, *X* at stationarity (that is *t* –→ ∞), as a function of both the bifurcation parameter *α* and noise intensity *σ*. In Figure 2.e–h we show the entropy change for the pitchfork bifurcation. Two observations are readily apparent: (i) as *α* is varied, entropy first increases and then decreases, at all noise levels considered here. Where this switch occurs depends on the level of noise: for no or low levels of noise, the switch happens at the bifurcation point, *α* = 0; for medium and large noise the happens well after the bifurcation point *(e.g*. *α* ≈ 2.5 for *σ* > 1); (ii) after the maximum in entropy has been reached, the decrease in entropy with increasing *α* happens at different speeds: rapidly for no noise; and much more slowly for noisy dynamics.

We consider the quasi-potentials for the transcritical and saddle node bifurcations in the appendix (Figure A1 and A2). Noise modifies the shape of the empirical quasipotential landscapes, just as in the case of the pitchfork bifurcation; see Figure A1 and A2, b-d). These variations become more pronounced with increasing noise (*i.e. σ*). For the transcritical and saddle node bifurcations, the behaviour of the entropy of the state variable *X*, however, is different as shown in the appendix, Figures A1 and A2, e-h. Neither do we observe distinct phases where entropy increases and decreases; nor is there a switch in entropy behaviour at the bifurcation point. In the case of the transcritical bifurcation (Figure A1, e-h), the entropy increases with *α*, followed by only a slight decrease in the slope after the *α* = 0; noise shifts this slightly. For the saddle node bifurcation (Figure A2, e-h) we observe that the entropy value decreases with *α*, followed again by a slight decrease in the slope after the bifurcation point, *α* = 0, but really only clearly discernible at medium and large noise.

The quasi-potentials corresponding to the different co-dimension 1 bifurcations clearly reflect the qualitative changes inherent to the deterministic dynamics. But for moderate and large (in the context of what we consider here) levels of noise we find that the interplay between noise and dynamics changes the overall qualitative behaviour.

### 3.2 Noise non-trivially increases variability in co-dimension 1 bifurcations

We consider three types of co-dimension 1 bifurcations which are the supercritical pitchfork bifurcation, the transcritical bifurcation and the saddle node bifurcation. The respective bifurcation diagrams are shown in Figure 3a.

**Figure 3:**
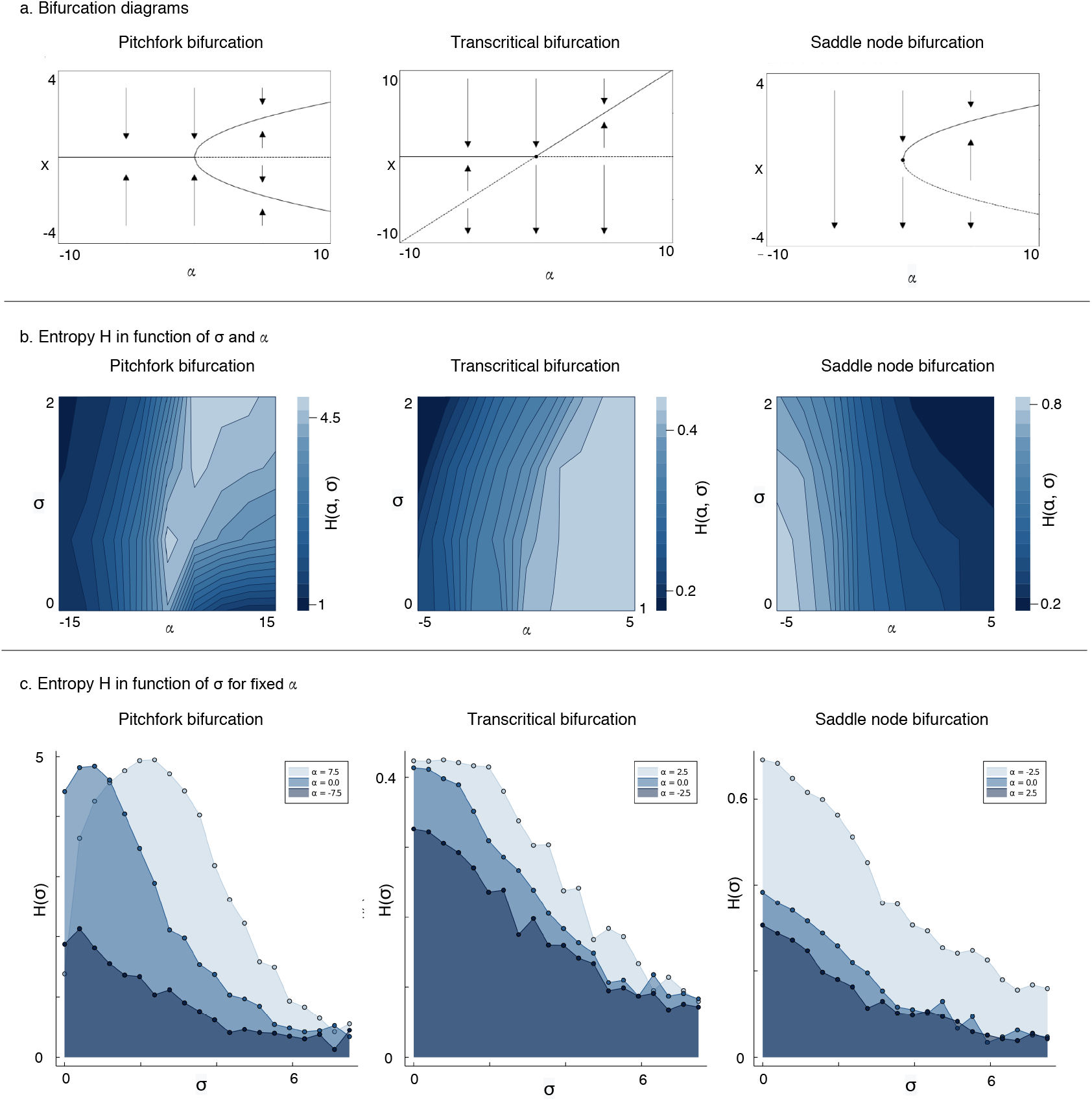
The entropy over the state space is shown for systems undergoing the supercritical pitchfork bifurcation, the transcritical bifurcation and the saddle node bifurcation (Equation 4–6). In the bifurcation diagrams (a) the stable fixed points and the unstable fixed points are indictaed by solid and dashed lines, respectively. Entropies values are displayed as functions of noise term σ (Equation 3) and the bifurcation parameter α (b). Additionally, the entropies are also shown as functions of σ for selected α values (–7.5,0.0 and 7.5 for the pitchfork bifurcation, and –2.5,0.0 and 2.5 for the transcritical and saddle node bifurcations).

Alongside the bifurcation diagrams in Figure 3a, we show contour plots of entropy values depending on the bifurcation parameter *α* and noise level *σ* in Figure 3b. In the case of the pitchfork bifurcation (Figure 3b, left), entropy increases progressively with *α* and uniformly across *σ* before then decreasing again. For low levels of noise, the maximal entropy is reached in the vicinity of the bifurcation point, *α* = 0. For positive α, entropy increases with noise, *σ*, over the ranges considered here. For the two other bifurcations, the contour plots are simpler. The transcritical bifurcation quasi-potential landscape (Figure 3b, center) displays a progressive increase in entropy as *α* increases. By contrast, for the saddle node bifurcation quasi-potential landscape (Figure 3b, right) the entropy of the state decreases as *α* increases, and it does so more rapidly as noise increases.

In Figure 3c we show the change in entropy with σ for three different values of the bifurcation parameter, *α* ∈ {-7.5,0, 7.5} for the pitchfork bifurcation, and *α* ∈ {-2.5,0,2.5} for the transcritical and saddle node bifurcations, respectively. For the pitchfork bifurcation we see that the entropy peaks for finite *σ* for all three *α* values considered. This suggests that at the transition-state of the dynamics (the vicinity of the bifurcation point), noise is modulated. In fact, for all three bifurcations the observations are similar: a decrease of entropy values with the increase of *σ*. However, the pitchfork bifurcation (Figure 3a, left) shows a transient increase for low values of noise, suggesting that here the noise non-trivially interacts with the dynamics especially around the transition state (the bifurcation point). Overall it appears that for fixed bifurcation parameter the entropy – quantifying the uncertainty about the state X of the system– ultimately to the level of noise. But this counter-intuitive result [22] can be explained by the way the the quasi-potential function changes shape, and ultimately flattens for large values of *σ*.

### 3.3 The relationship between empirical quasi-potential function and the potential function

In order understand the behaviour of the entropy, we need to compare the quasipotential function, *Q*(*X*), given by Eqn. (7), with the potential function, *U*(*X*), which is here used to define the gradient system. In Figure 4, we contrast the potential functions (a-d) and the corresponding empirical quasi-potentials (e-h). For geometric random noise the positioning of the valley bottoms at *X* = 0 for negative *α* and at 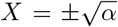 for positive *α* remain unchanged (XXX). Thus we expect the dynamics of the stochastic system to be centered around the stationary points of the deterministic system. There are, however, also striking differences: (i) in the potential surface we observe smoother, more gradual changes in the landscape, whereas the valleys of the quasi-potential landscape are much more localised; (ii) the unstable fixed point at *x* = 0 for positive *α* is discernible in the potential but not in the quasi-potential surface. This is not surprising, as the system will never attain this state as *t* —→ ∞.

**Figure 4:**
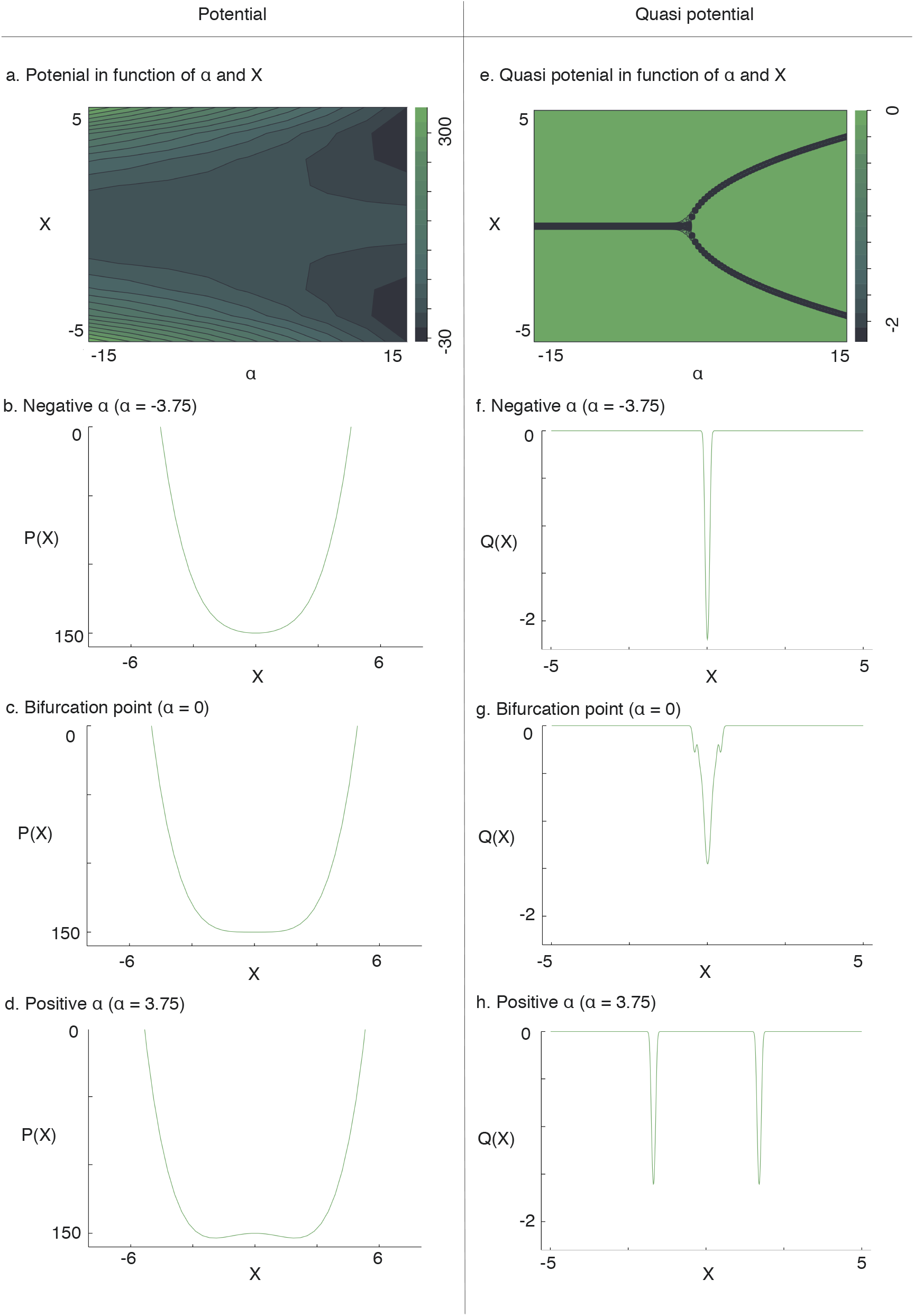
Potential function P (X) (a-d) and quasi-potential function Q(X) (e-h)for a deterministic system undergoing the supercritical pitchfork bifurcation (Equation 4). The example presented includes a two dimensional comparison of P (X) and Q(X) in function of the bifurcation parameter α and the state variable X (a and e) as well as one dimensional comparisons of P(X) and Q(X) for three selected values of α which represent the three qualitatively distinct global stability outcomes of the system whilst undergoing the bifurcation. They are before the bifurcation event, α = −3.75 (b,f); at the bifurcation event, α = 0.0 (c, g); and after the bifurcation event, α = 3.75 (d, h).

We note that the empirical landscapes was, for practical purposes, calculated for large but finite times. In the large time limit, the probability distribution for the deterministic system (*σ* = 0) would be a Dirac *δ*-function and the landscape then becomes -*δ*(*X*) (as can be seen by taking the limit of a suitable approximation for *δ* and substituting it into Eqn. (7)). Because we only simulate for finite times, our representation for *σ* = 0 does not show this extreme shape. In Figure A3 in the appendix we illustrate the effects of finite time scales numerically.

## 4 Discussion

The objective of this study has been to analyse systematically the effects of increasing a geometric random noise on dynamical systems undergoing co-dimension 1 bifurcations that have been implicated in cell-fate decision making.

We have quantified uncertainty of the dynamical system undergoing three different bifurcations (pitchfork, transcritical and saddle node) via the entropy. A number of studies have considered such systems as frameworks for interpreting results from single cell transcriptomic analysis. For instance, a pitchfork bifurcation can be seen as a differentiation into two different lineages [2,12]. We find an increase in entropy with a peak at the bifurcation point, but this becomes less pronounced as the noise intensity increases. The “rise and fall shape” of uncertainty across transition processes such as differentiation (into two lineages, reproducing a pitchfork bifurcation) has been observed in several studies (using mainly but not only entropy as an uncertainty measure) [23–25].

Biologically the increase in entropy may be relevant for the induction phase of cellular differentiation to promote the transition from one valley (i.e. cell type) to another. The decreasing uncertainty may be reflected in the stabilisation phase of the system after the bifurcation point [26]. Our results suggest that this transient elevation of uncertainty is inherent to bifurcation processes. In our analysis simulations we also find that the switch point between the increasing and decreasing phases shifts as noise increases; transcriptional variability inherent in cellular dynamics may therefore directly (and endogenously) influence the modality of the cell fate decision making process [26,27].

For the two other bifurcations, the transcritical and the saddle node bifurcations, this rise and fall shape is not observed. Compared to the pitchfork bifurcation both systems only have one stable fixed point: for all values of α in the transcritical bifurcation case and for positive values of *α* in the saddle node case (see Figure 3, top row). First, the transcritical bifurcation can be interpreted as cells transitioning from one stable state to a different state with the branching event inducing a switch in the states’ stabilities. Thus, the first stable state becomes an unstable state after the branching event, whereas the unstable state becomes the new stable state at the bifurcation. Second, the saddle node bifurcation has been interpreted as a transition between cell states where one state is destroyed while a new is created [12]. Like the transcritical bifurcation there is only one stable fixed point for all values of the bifurcation parameter, but there is no conversion between stable and unstable fixed points; the hills and valleys in the landscape are simply destroyed at the (deterministic) bifurcation.

A biological example suitable for the transcritical or saddle node bifurcation could be a single lineage differentiating, such as the *in vitro* avian erythroid differentiation [28]. In this model, T2EC progenitor cells can only differentiate into one fate [29]. The question then arises whether the transcritical or the saddle node suits the best to describe this differentiation. Using single cell analysis during this erythropoiesis process, rise and fall shape of the uncertainty has been observed [30]. However, we do not observe this variation in our dynamical systems undergoing these bifurcations. Indeed, entropy increases along α for the transcritical bifurcation and decreases for the saddle node bifurcation, without any switch around the branching event in both cases. In addition, the addition of noise slightly changes the shape but without net effects compared to the pitchfork case. Bifurcations that allow for multistability such as pitchfork bifurcation in our study matches well with biological transition processes (branching shape, entropy and effect of noise) whereas single fixed point bifurcations remain difficult to reconcile with some experimental observations. The reasons may originate from the fact that *in vivo* monolineage differentiation *(i.e.* unipotency) is often imposed on precursor cells, which have low differentiation potential and are not well represented in literature [31]. In that case, we simply have a lack of data concerning these type of differentiation in order to draw firm conclusions about the qualitative dynamics in such systems. It is also important to realise that multistability can arise in different ways; not all of these are necessarily apparent at the transcriptomic level. For example, shuttling of transcription factors between nucleus and cytoplasm can give rise to multistability [32], and has been observed experimentally [33].

All together, these results suggest that the change in entropy across cell differentiation depends of the type of the transition (single-fate or multi-fate) and is sensitive to the noise levels [34]. Our analysis, taken together with experimental observations, suggests that a much more nuanced consideration of the potential differentiation dynamics and noise may be advisable. For all three bifurcations, we observe a decrease of the entropy at specific bifurcation points (3 different values of *α*) with the increase of noise strength. There appears to be a tension between the change in entropy as the bifurcation parameter is varied, and the decrease in entropy as noise-levels increase. Indeed, in parallel with the transient increase of entropy observed during differentiation processes, an apparently counter-intuitive increase of gene-to-gene correlation has been reported [30,35]. It is known that noise and dynamics can give rise to complex behaviour of entropy and related measures [36]. Changes in entropy across differentiation have been observed (experimentally and theoretically) [37,38]. But concomitantly we can also often see an increase of gene-to-gene correlations, which could be a direct consequence of the external signals received to differentiate, activating genes and features allowing cells to achieve a reproductive transition [29]. Noise is probably allowing genes to explore the state space a more widely and this may be ultimately linked to induction of cell-fate transitions.

Waddington’s epigenetic landscapes have influenced our way to see and interpret cell dynamics across developmental processes; we show that there are systematic differences between the potential (depicted by Waddington) and the empirical or quasi-potential landscapes: the quasi potential is much more localised than the potential: the system’s dynamics, even with noise present explore only the valley bottoms of the potential landscape.We note that this may entail practical limitations for reconstructing the (deterministic) dynamics from single cell data (or, as here, solutions of the stochastic dynamics.

In order to gain clearer insights into the mechanisms underlying cell fate decision-making, we believe that more detailed analyses of bifurcations – qualitative changes in system dynamics – will require further analysis: if even simple co-dimension 1 bifurcations can exhibit such rich dynamics, then more complicated bifurcation systems will require much more detailed and careful analysis. Purely data-driven analysis of such systems may otherwise risk missing or misinterpreting signals in single cell data.

## Acknowledgments

We would like to thank the members of the Theoretical Systems Biology Group for many fruitful discussions.

## A Appendix

### A.1 Changing landscapes for the transcritical bifurcation

**Figure A1:**
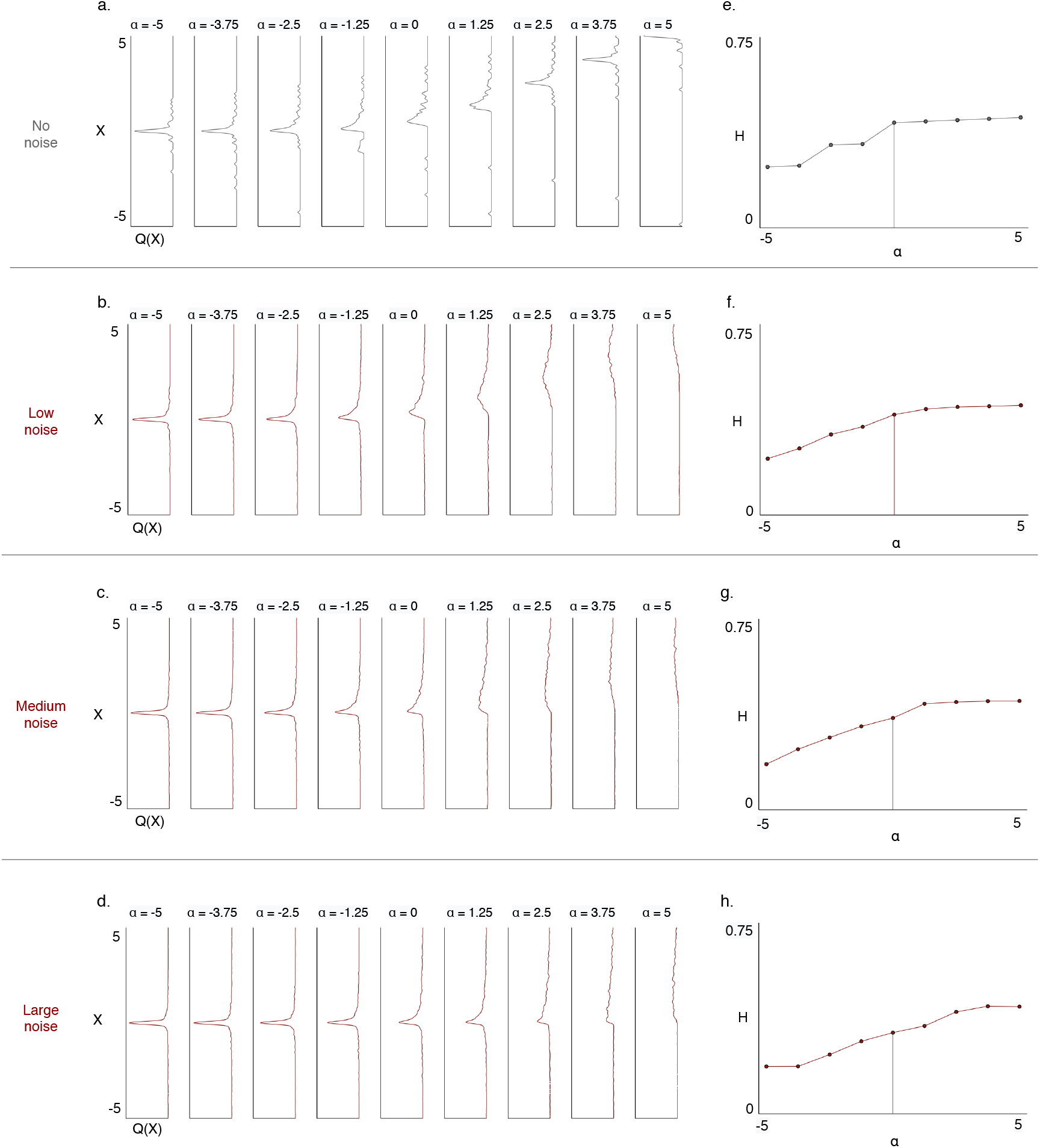
The quasi-potential (a-d) and the entropy over the state space (e-h) are displayed as the system undergoes the transcritical bifurcation (Equation 5) for the deterministic case (a, e) and three selected stochastic cases (b, c, d and f, g, h). The noise levels are σ = 0.0, σ = 0.6, σ = 1.3, and σ =1.6 where σ is the scaling factor in the noise term in Equation 2.

### A.2 Changing landscapes for the saddle node bifurcation

**Figure A2:**
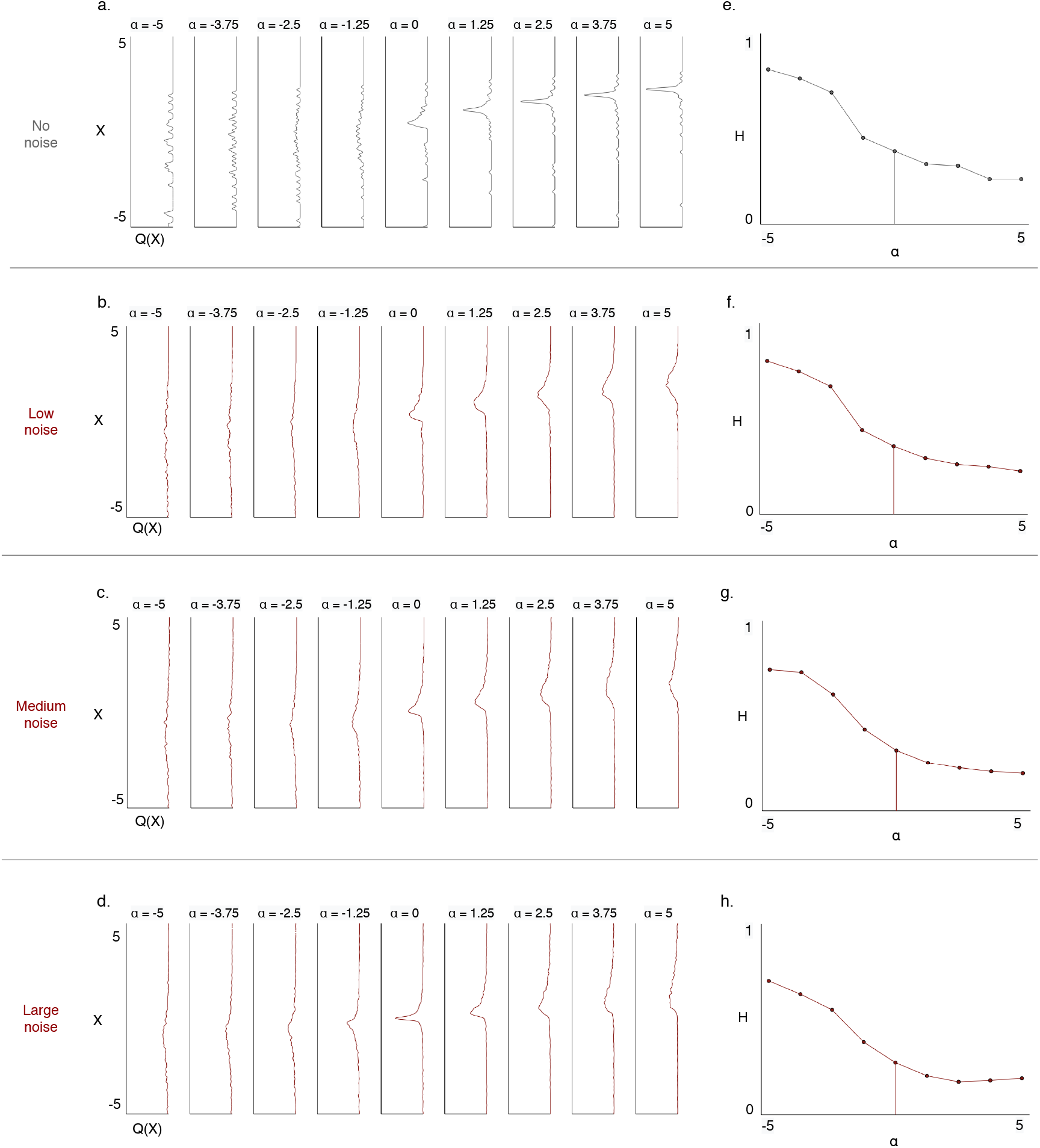
The quasi-potential (a-d) and the entropy over the state space (e-h) are shown as the system undergoes the saddle node bifurcation (Equation 6). The four different levels of noise are σ = 0, σ = 0.6, σ = 1.3, and σ = 1.6 (top to bottom row, respectively) where σ is the scaling factor in the noise in Equation 2.

### A.3 Numerical Considerations

We explored different simulation settings in order to understand how they effect the shape of the retrieved quasi-potential surface for the supercritical pitchfork bifurcation (as a function of the bifurcation parameter *α* and the state of the system X). The time span and the numbers of SDE solutions clearly limit the insights we can gain into the shape of the quasi-potential (see Section 2.2 and Figure 1 for details on the workflow). All calculations presented in the main part of the manuscript were carried out with the final parameter setting.

**Figure A3:**
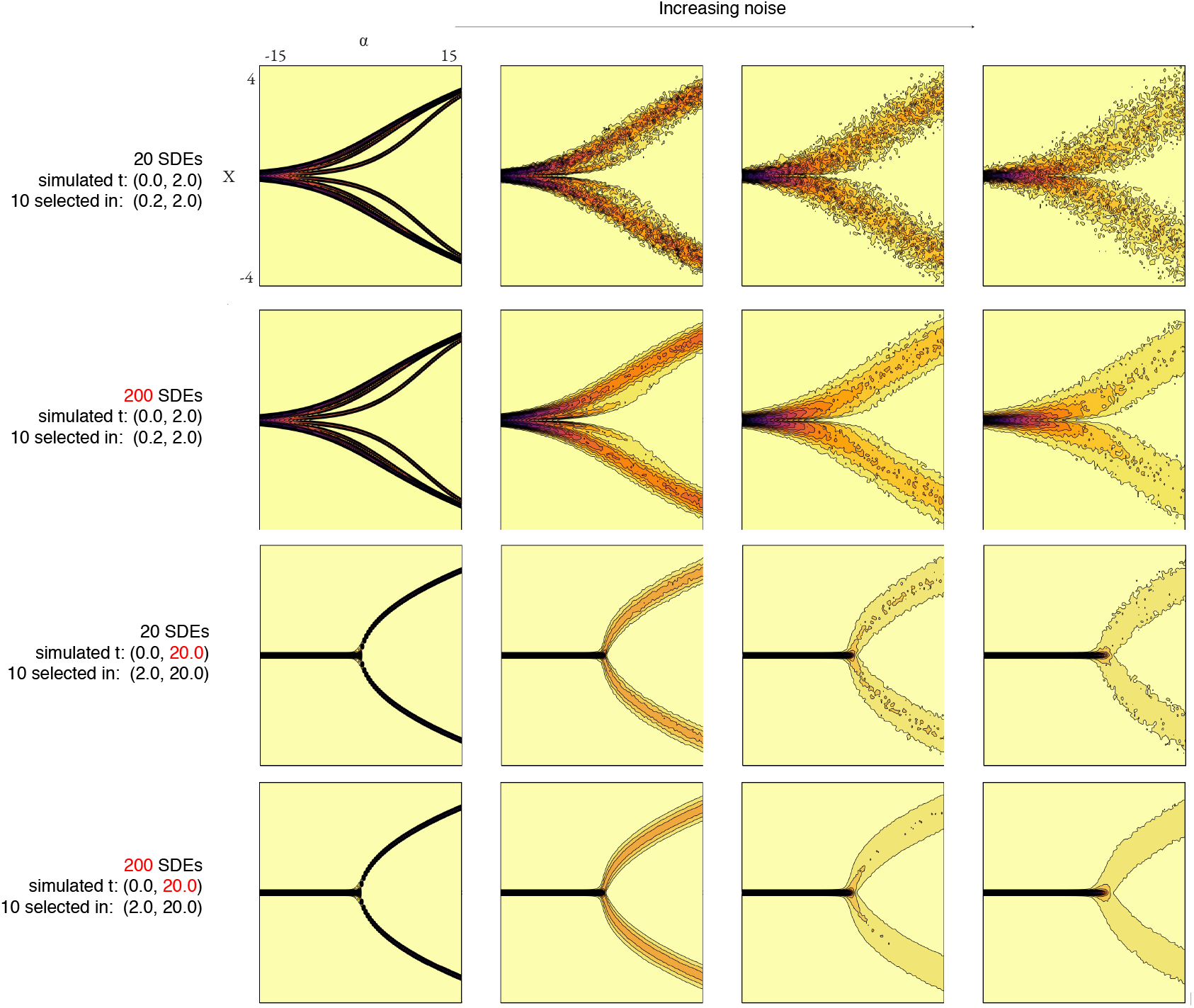
The quasi-potential surfaces for four different settings of the empirical quasi-potential calculation are displayed. The quasi-potential surface is shown for four different settings (row) and four different levels of noise (column). From the left to the right, the four different levels of noise are α ∈ (0,0.6,1.3,1.6). The settings vary in the number of SDE solutions taken into account for each initial condition, and the time span of each simulation.

